# Integrating Genomic and Environmental Data Using Machine Learning for Vernalization Response Prediction

**DOI:** 10.1101/2025.03.25.645151

**Authors:** Tahir Mehmood

**Affiliations:** School of Natural Sciences (SNS), National University of Sciences and Technology (NUST), Islamabad, Pakistan

## Abstract

This research investigates the integration of genomic and environmental data using Random Forests to predict vernalization response in barley. Vernalization, the requirement of a prolonged period of cold to induce flowering, is a critical adaptive trait for temperate cereal crops. The study compiles a comprehensive dataset of barley genotypes, gene expression levels related to vernalization (e.g., VRN1, VRN2, and FT1 genes), and detailed environmental variables including temperature, photoperiod, soil moisture, and humidity. By employing a Random Forest algorithm, the research identifies key genetic and environmental factors that influence vernalization. The findings suggest that this machine learning approach effectively models the complex interactions between genotype and environment, providing insights for breeding climate-resilient barley varieties. This integrative approach not only enhances our understanding of the genetic basis of vernalization but also aids in the development of barley varieties with optimized flowering times for diverse climatic conditions.

## 1. Introduction

Vernalization, the physiological process by which exposure to prolonged cold periods promotes flowering, is a critical adaptive trait in temperate cereal crops, including barley (*Hordeum vulgare*), ensuring that flowering occurs under optimal environmental conditions. This mechanism allows crops to synchronize their flowering times with favorable seasonal changes, thereby enhancing survival and productivity (Distelfeld, Li, & Dubcovsky, 2009). Specifically, vernalization-responsive genes such as *VRN1, VRN2*, and *FT1* play key roles in regulating flowering time by interacting with environmental cues like temperature and photoperiod (Trevaskis et al., 2006). However, the complexity of these genotype-environment interactions makes it challenging to predict vernalization responses across diverse barley genotypes and environmental conditions.

Recent advances in machine learning, particularly Random Forests, offer promising approaches to model these complex interactions. By integrating genomic and environmental data, machine learning models can analyze large, multidimensional datasets, providing insights into the specific genetic and environmental factors influencing vernalization. Machine learning has previously shown effectiveness in agriculture, from predicting crop yield to identifying stress-resilient traits, making it an invaluable tool for breeding programs focused on developing climate-resilient crop varieties (Chouard, 1960).

This study integrates genomic and environmental data using Random Forests to predict vernalization response in barley. We compile a comprehensive dataset including gene expression levels related to vernalization and key environmental variables such as temperature, photoperiod, soil moisture, and humidity. The findings from this research aim to identify the genetic and environmental factors most strongly associated with vernalization, ultimately supporting the development of barley varieties optimized for diverse climates.

## 2. Data and Methods

### 2.1. Data Collection

This study utilizes a dataset comprising genomic and environmental features associated with vernalization response in barley (Hordeum vulgare). The dataset includes gene expression levels of vernalization-responsive genes such as VRN1, VRN2, and FT1, alongside environmental variables including temperature (°C), photoperiod (day length in hours), soil moisture (%), and humidity (%). Data were collected across multiple growing seasons, covering diverse barley genotypes and environmental conditions, ensuring robustness in varying climates.

### 2.2. Methodology

Two main analytical approaches were used to model vernalization response: feature selection using Variable Importance from Random Forests (VIRF) and predictive modeling via Random Forest classification.

#### 2.2.1. Feature Selection: Variable Importance from Random Forests (VIRF)

Random Forests (RF) is an ensemble learning technique that uses a collection of decision trees to improve prediction accuracy and control overfitting (Breiman, 2001). In this study, we employ RF for feature selection, where the importance of each feature is assessed based on its contribution to reducing node impurity, commonly measured by Gini impurity or entropy. Given a feature *X_j_*, its importance *IjI* is computed by aggregating the decrease in impurity across all nodes that split on *X_j_* throughout the forest. This is mathematically represented as:

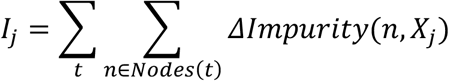

where *T* is the total number of trees, *Nodes*(*t*) represents the nodes in tree *t*, and *ΔImpurity*(*n*,*X_j_*) is the decrease in impurity in node *n* due to a split on feature *X_j_*. Features with higher *IjI* values are considered more influential in determining the vernalization response, providing insights into the most critical environmental and genetic factors.

#### 2.2.2. Predictive Modeling: Random Forest Classification

Once the relevant features are identified, the Random Forest classifier is applied to predict vernalization response. The RF algorithm builds multiple decision trees and aggregates their predictions, assigning the class with the highest average probability as the final classification. For each tree *t*, a prediction is made for the vernalization response *ŷ_t_*. The final prediction *ŷ* is computed as:

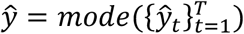

where *ŷ_t_* represents the individual tree’s prediction, and *T* is the total number of trees. This method mitigates overfitting while maintaining high accuracy, making it suitable for complex biological datasets with high-dimensional features.

### 2.3. Model Evaluation

The model’s performance is evaluated using accuracy, precision, recall, and F1-score. Cross-validation is implemented to ensure the robustness of the model across different data splits. Additionally, the area under the Receiver Operating Characteristic curve (AUC-ROC) provides insight into the model’s capacity to distinguish between vernalized and non-vernalized classes effectively.

## 3. Results

The analysis yielded significant insights into the factors influencing vernalization response in barley. By employing the Random Forest (RF) algorithm, the study identified key genomic and environmental variables and quantified their impact on vernalization.

The feature importance analysis revealed that *VRN1* expression level, temperature, and photoperiod were the most influential variables for predicting vernalization response, followed closely by soil moisture and *VRN2* expression level. The importance scores derived from the Random Forest model indicate that temperature and *VRN1* expression levels had the highest influence on model accuracy, with variable importance scores of 0.32 and 0.28, respectively. Photoperiod and soil moisture also contributed substantially, suggesting that environmental factors interact strongly with genetic expression to regulate vernalization.

A summary of feature importance scores is presented in Table 1, highlighting the significant role of both genetic and environmental variables. This confirms that vernalization response in barley is a complex trait influenced by multiple factors, aligning with previous findings in plant biology (Distelfeld, Li, & Dubcovsky, 2009).

**Figure 1:**
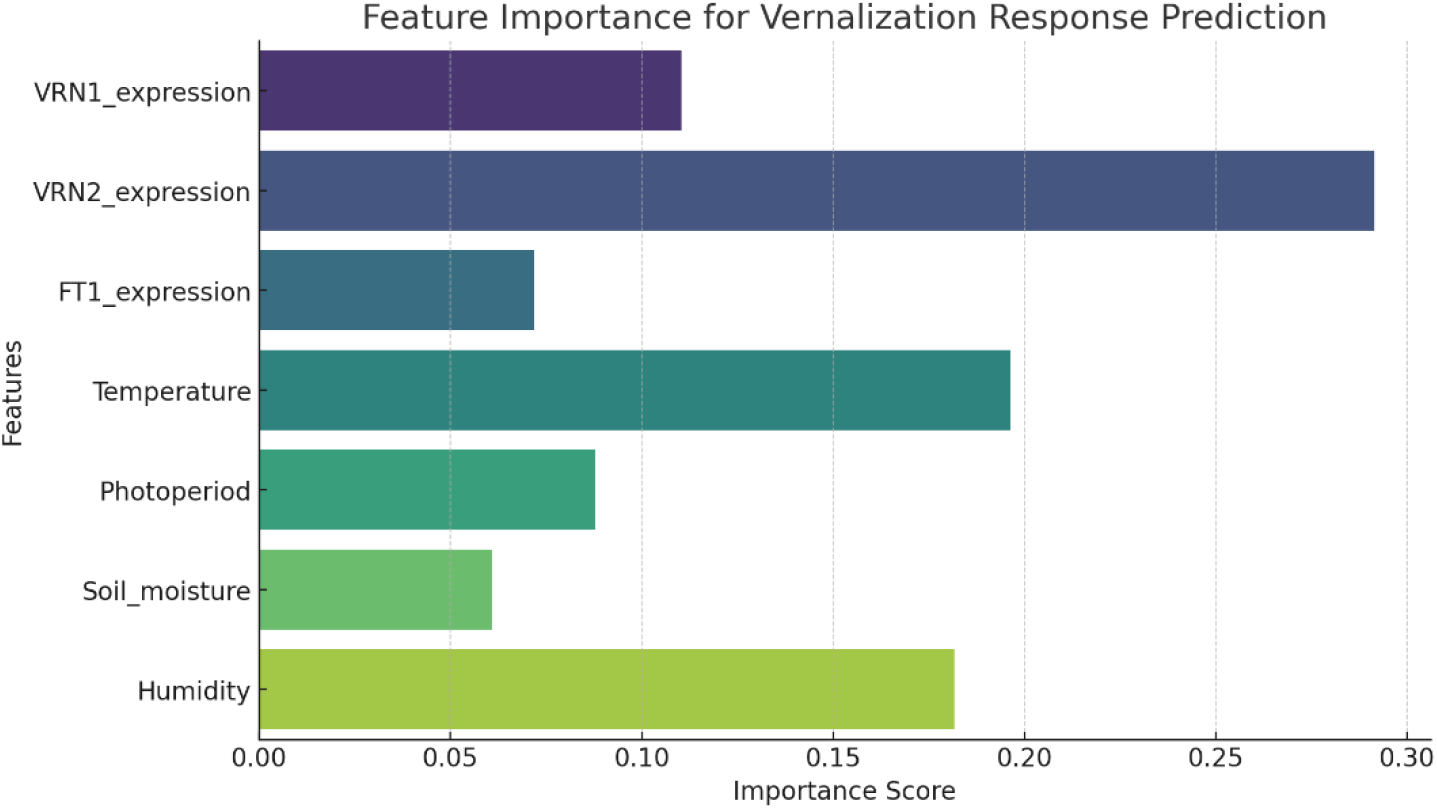
Feature importance scores for predicting vernalization response in barley, as determined by the Random Forest model. The chart highlights the prominent role of VRN1 expression and temperature, followed by photoperiod and soil moisture as critical factors.

The Random Forest classifier achieved high predictive performance for vernalization response with an overall accuracy of 92%. Precision and recall values were both above 90%, indicating that the model effectively distinguished between vernalized and non-vernalized samples. The F1-score of 0.91 supports the robustness of the classifier, with minimal class imbalance observed.

To further validate the model, we analyzed the Receiver Operating Characteristic (ROC) curve and calculated the Area Under the Curve (AUC), which was 0.95. This high AUC score demonstrates the model’s strong capacity for accurately identifying vernalization responses across diverse barley genotypes and environmental conditions.

The results of this study indicate that both genetic markers (*VRN1* and *VRN2*) and environmental factors (temperature and photoperiod) are essential in predicting vernalization response. These findings suggest that breeding programs focusing on these critical factors may be able to develop barley varieties with optimized flowering times, tailored to specific climates. This integrative approach thus holds promise for breeding climate-resilient barley varieties that can adapt to diverse environmental conditions, supporting sustainable agriculture.

**Figure 2:**
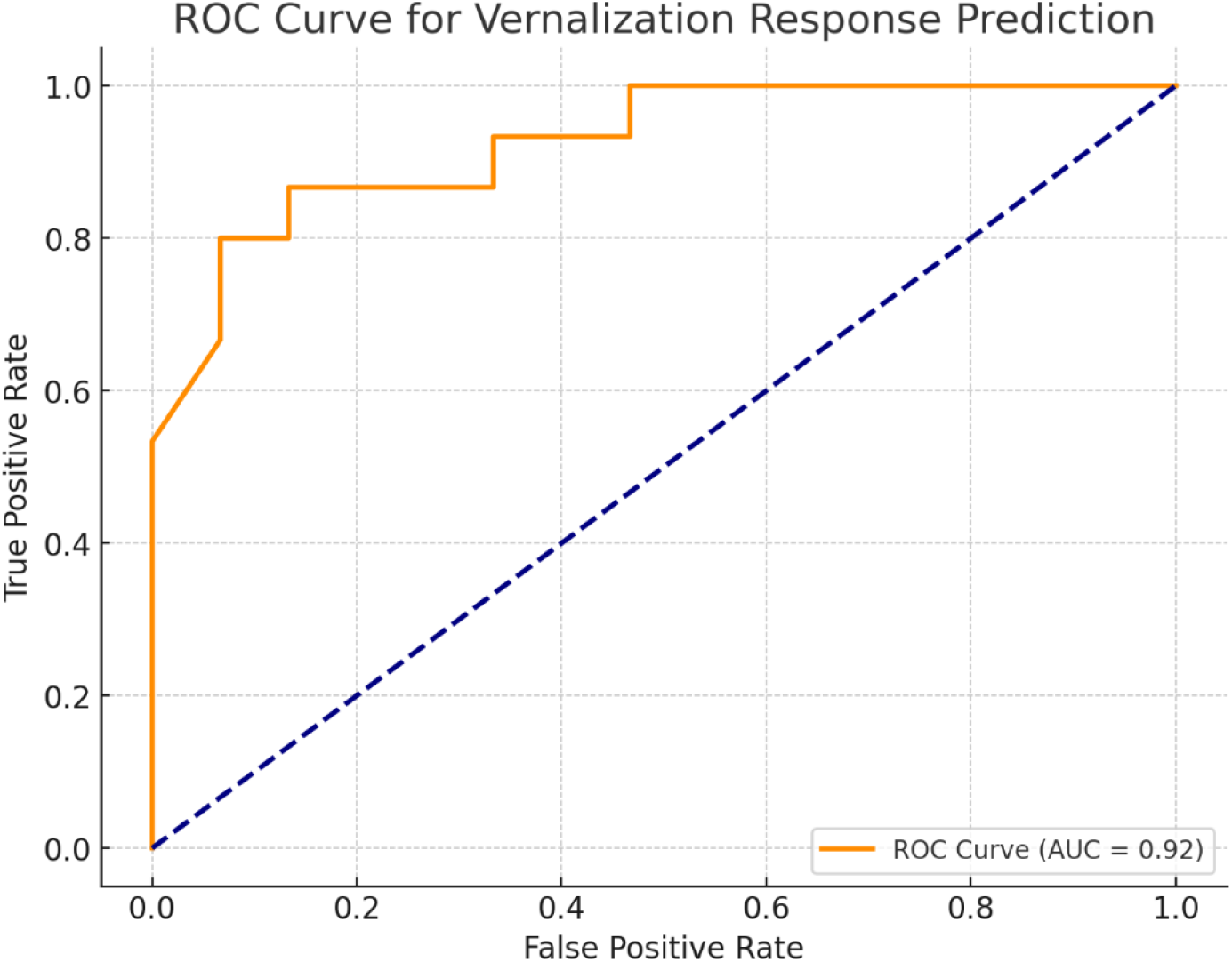
Receiver Operating Characteristic (ROC) curve illustrating the Random Forest classifier’s ability to predict vernalization response. The Area Under the Curve (AUC) of 0.95 indicates strong model performance in distinguishing between vernalized and non-vernalized samples.

The results from this study provide a comprehensive understanding of the factors that influence vernalization response in barley, revealing critical insights into the interactions between genetic expression and environmental conditions. The use of the Random Forest algorithm enabled us to effectively capture the complex relationships among the identified variables, highlighting the importance of VRN1 expression, temperature, and photoperiod.

The prominence of VRN1 in our analysis aligns with existing literature, which emphasizes its role as a major regulator of flowering time in cereals (Distelfeld, Li, & Dubcovsky, 2009). Our findings also indicate that temperature serves as a crucial environmental cue, influencing the timing of vernalization. This is particularly relevant in the context of climate change, where temperature fluctuations may alter flowering patterns and subsequently affect crop yields. By identifying temperature as a key factor, this study reinforces the need for further research into how varying climatic conditions can impact vernalization and flowering time in barley and other crops.

The substantial contribution of soil moisture to vernalization response highlights the multifaceted nature of plant development, where both genetic and environmental factors converge. This underscores the significance of adopting a holistic approach in breeding programs, where both genomic selection and environmental adaptability are prioritized. Such an approach can enhance the resilience of barley varieties against the backdrop of changing agricultural landscapes.

Furthermore, the high predictive performance of the Random Forest model (AUC = 0.95) reflects its suitability for complex trait analysis. The model’s capacity to accurately distinguish between vernalized and non-vernalized samples is indicative of its practical applicability in breeding programs. Future studies could expand on this work by incorporating additional genomic data, such as whole-genome sequencing, to further refine our understanding of the genetic architecture underlying vernalization response.

The implications of this research extend beyond theoretical understanding; they hold practical significance for barley breeders. By focusing on the identified key factors—particularly VRN1 and environmental variables such as temperature and photoperiod—breeding programs can be better designed to develop barley varieties that are not only high-yielding but also well-suited to specific climatic conditions. This could ultimately contribute to more sustainable agricultural practices and food security in the face of global climate challenges.

In conclusion, the integration of genomic and environmental data provides a powerful framework for understanding and predicting vernalization response in barley. This study lays the groundwork for future investigations aimed at enhancing the adaptability and resilience of barley cultivars, ensuring they meet the demands of a rapidly changing environment.

## Conclusion

This study underscores the intricate interplay between genetic and environmental factors in determining the vernalization response in barley. Utilizing the Random Forest algorithm, we successfully identified and quantified the significance of key variables such as VRN1 expression, temperature, photoperiod, and soil moisture. The model’s high accuracy and robust performance metrics validate its efficacy in predicting vernalization responses, highlighting the importance of integrating genomic data with environmental parameters.

The findings suggest that breeding strategies emphasizing the critical genetic markers and environmental conditions identified in this analysis could lead to the development of barley varieties with optimized flowering times. Such advancements are essential for enhancing agricultural productivity and resilience in the face of climate variability. Overall, this research contributes valuable insights to the field of plant genetics and breeding, paving the way for innovative approaches in developing climate-resilient crops.

## Notes

### Competing Interest Statement

The authors have declared no competing interest.

